# ISME - Incoherent Sampling of Multi-Echo data to minimize cardiac-induced noise in brain maps of R_2_* and magnetic susceptibility

**DOI:** 10.1101/2025.02.05.636576

**Authors:** Quentin Raynaud, Rita Oliveira, Nadège Corbin, Yaël Balbastre, Ruud B. van Heeswijk, Antoine Lutti

**Author notes:** Correspondence Antoine Lutti Laboratory for Research in Neuroimaging, Department for Clinical Neuroscience, Lausanne University Hospital, Ch. de Mont-Paisible 16, CH-1011 Lausanne. These authors contributed equally to this work (Quentin Raynaud, Rita Oliveira).

## Abstract

**Purpose:** Maps of the MRI parameters R2* and magnetic susceptibility (𝜒) enable the investigation of microscopic tissue changes in brain disease. However, cardiac-induced signal instabilities increase the variability of brain maps of R2* and 𝜒. In this study, we introduce ISME – a sampling strategy that minimizes the level of cardiac-induced instabilities in brain maps of R2* and 𝜒.

**Methods:** ISME uses phase-encoding gradients to shift the k-space frequency of the acquired data between consecutive readouts of a multi-echo train. As a result, the multi-echo data at a given k-space index is acquired at different phases of the cardiac cycle. We compare the variability of R2* and 𝜒 maps acquired with ISME and with standard multi-echo trajectories in N=10 healthy volunteers. We investigate the effect of both trajectories on the spatial aliasing of pulsating MR signals and propose a weighted-least squares (NWLS) approach for the estimation of R2* that accounts for the increase of the residuals with echo time.

**Results:** ISME reduces the variability of R2* and 𝜒 maps across repetitions by 25/26/21% and 24/32/23% in the cerebellum/brainstem/whole brain, respectively. With ISME, the spatial aliasing of pulsating MR signals is incoherent between raw echo images, leading to visually sharper R2* maps. The proposed NWLS approach for the estimation of R2* reduces the dependence of the fitting residuals on echo time and the variability of R2* by an additional 3/2/1% in the cerebellum/brainstem/whole brain.

**Conclusion:** ISME allows the mitigation of cardiac-induced signal instabilities in brain maps of R2* and 𝜒, improving reproducibility.

## 1. Introduction

The transverse relaxation rate (R2* = 1/T2*)^1^ and magnetic susceptibility (𝜒)^2^ are markers of iron and myelin content within brain tissue^3,4^ and allow the study of microscopic tissue change due to brain disease in patient populations.^5–7^ Maximal reproducibility is required of R2* and 𝜒 maps to allow optimal sensitivity to brain change in neuroscience studies. In addition to thermal noise,^8,9^ MRI data reproducibility is affected by signal instabilities of physiological origin.^10,11^ Respiration and cardiac pulsation are the primary sources of physiological signal instabilities, which can also cause image aliasing or blurring.^12,13^ Physiological signal instabilities increase with the echo time used for data acquisition.^14,15^ Gradient-recalled echo (GRE) data used for R2* and 𝜒 mapping typically reach echo times of up to ∼40ms^7,16,17^ and are therefore particularly sensitive to physiological signal instabilities.^16,18^

Respiration-induced signal instabilities originates from periodic fluctuations of the magnetic field due to changes in air volume in the lungs.^19^ The effects of respiration on brain MRI data are typically corrected using phase navigators that reduce the variability of R2* and 𝜒 maps by 23% and 12% across repetitions in white matter.^16^ Cardiac-induced signal instabilities are the result of multiple physiological mechanisms: cardiac pulsation generates a pressure wave that leads to head motion,^20^ CSF pulsation^21,22^ and deformation^23,24^ of inferior regions such as the brainstem^25,26^ and cerebellum.^27^ Also, pulsatile vessel movement and periodic blood influx^28–32^ lead to spatially-localized effects that are most pronounced in highly vascularized regions such as the orbitofrontal cortex^22,33^ and periventricular regions.^33,34^ Physiological effects that are coherent across a voxel, such as variation of the mean magnetic field across a voxel,^16,35^ laminar flow,^36,37^ and motion^38–40^ result in a net phase shift of the signal and primarily affect magnetic susceptibility estimates. On the other hand, effects that are incoherent across a voxel (e.g. turbulent flow,^41^ local B0 inhomogeneities^42^) result in a net change of the magnitude of the signal and primarily affect R2* estimates. Cardiac pulsation has recently been reported to lead to variations of R2* by up to 3s^-1^ across the cardiac cycle and to account for ∼35% of the variability of R2* maps across repetitions.^18^

The acquisition of R2* and 𝜒 data stretches over several minutes. Cardiac-induced signal instabilities cannot be regressed out retrospectively from the temporal evolution of the signal as is routinely done for functional MRI data.^12,43,44^ Also, the effects of cardiac pulsation on the MR signal are spatially localized and are not compatible with the navigator techniques used for the correction of respiration- or motion-induced effects.^45,46^ Instead, prospective strategies show promising potential to reduce the impact of cardiac-induced signal instabilities in the acquired data. Specially-designed k-space sampling trajectories lead to spatially-incoherent aliasing of cardiac-induced signal instabilities across the field of view of an individual image.^47,48^ Setting the number of samples at each k-space index according to the local amplitude of the signal instabilities is effective to improve the reproducibility of R2*-mapping data.^49^ However it does not consider the effect of cardiac-induced signal instabilities across the temporal dimension of the multi-echo data.

In this study, we introduce a new sampling strategy to reduce the level of cardiac-induced signal instabilities in multi-echo data used for the computation of R2* and 𝜒 maps. This sampling strategy is based on dispersing the effect of the instabilities along the temporal dimension of the multi-echo data: with ISME (Incoherent Sampling of Multi-Echo data), the k-space frequency of the acquired data is changed between consecutive readouts of the echo train. As a result, the multi-echo data at a given k-space index are acquired across different phases of the cardiac cycle. The goal of this study was to demonstrate that ISME leads to higher reproducibility of brain maps of R2* and 𝜒 than standard k-space sampling trajectories in healthy volunteers. We also investigated the effect of both sampling strategies on the spatial aliasing of cardiac pulsation effects in maps of R2* and 𝜒 and on the fitting residuals.

## 2. Methods

### 2.1. ISME

With standard multi-echo trajectories, data are acquired at a single k-space frequency after each radiofrequency (RF) excitation. To allow for full image encoding, this process is repeated across a suitable set of k-space frequencies. Because the repetition time is much smaller than the cardiac period (TR ≲ 50ms), the multi-echo data at each k-space frequency can be assumed to have been acquired at the same phase of the cardiac cycle (Figure 1A). Because cardiac pulsation leads to systematic changes of R2* across the cardiac cycle,^18^ this multi-echo data contains exponential-like effects of cardiac pulsation on signal decay, leading to a bias of the apparent R2*. This bias depends on the phase of the cardiac cycle at the time of acquisition of the data (Figure 1B). To minimize the effect of cardiac pulsation on R2* estimates, we introduce ISME (Incoherent Sampling of Multi-Echo data) - a new sampling strategy that allows to sample the cardiac phase incoherently along the echo dimension. With ISME, short gradient pulses are played out between each readout of the multi-echo train to shift the k-space index (Figure 1C). Upon completion of data acquisition, the data at each k-space index is made up of multiple echoes acquired at different phases of the cardiac cycle. With ISME, estimation of R2* averages out the fluctuations of signal decay across the cardiac cycle, at the cost of larger fit residuals (Figure 1D).

**Figure 1:**
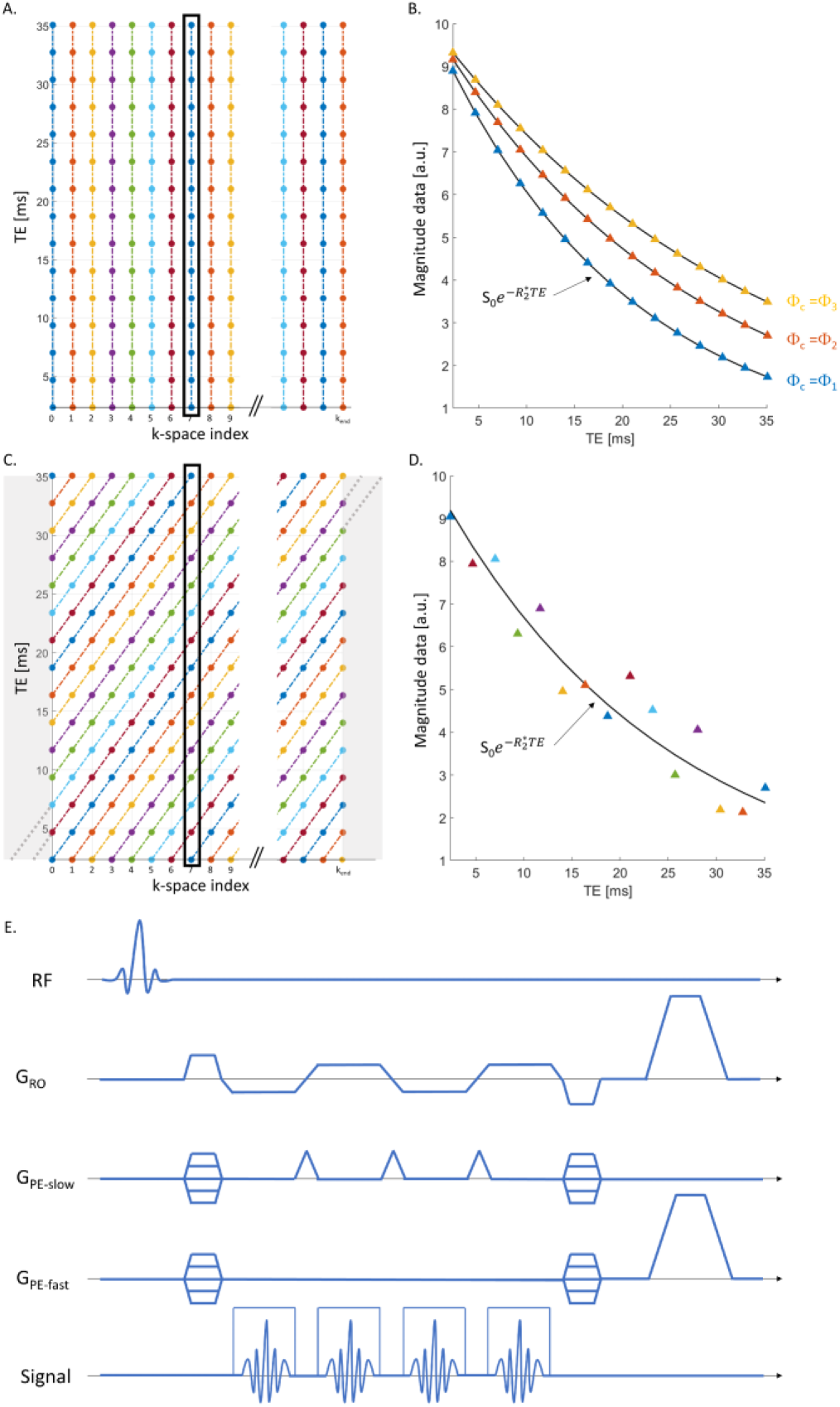
Traversal of the k-TE space with a standard multi-echo trajectory and with ISME. (A) The different colours represent different phases of the cardiac cycle. With a standard multi-echo trajectory, multi-echo data is acquired at a distinct k-space frequency following each radio- frequency excitation. Because the repetition time is much smaller than the cardiac period (TR ≲ 50ms), the multi-echo data at each k-space frequency is effectively acquired at the same phase of the cardiac cycle. (B) This multi-echo data contains exponential-like effects of cardiac pulsation on signal decay that depend on the phase of the cardiac cycle at the time of acquisition of the data. (C) To minimize the effect of cardiac pulsation on R2* estimates, the proposed ISME strategy shifts the current value of the k-space index between the readouts of the multi-echo train. (D) As a result, the multi-echo data of each k-space location is made up of points acquired at different phases of the cardiac cycle. (E) Pulse diagram of the ISME GRE sequence. With ISME, the shift of the k-space index is implemented by means of short gradient pulses along the slow phase-encode direction (PE), as in EPI. However, unlike EPI, the data from the consecutive readouts of the multi-echo train are stored separately to obtain an individual image at each echo time.

In this conceptual description of ISME, the k-space index is a simple counter that reflects the sequential ordering of data sampling. It is independent of the physical coordinates of the k-space frequencies and of the k-space trajectory used for data acquisition (e.g. linear, spiral or radial). Also, the time span between consecutive indexes might be a single TR interval in the case of two data points acquired consecutively, or a longer time interval. Here, we consider the case of a 3D FLASH sequence with linear sampling, in which neighbouring points are acquired consecutively along the fast phase-encoding direction, for each phase-encoding step along the slow direction. With this trajectory, the time interval between consecutive data points along the fast phase-encoding direction is TR ≲ 50ms, much shorter than the typical cardiac period. Cardiac pulsation leads to a broad modulation of the MR signal across k-space along this phase-encoding direction. We therefore implemented ISME along the slow phase-encoding direction, for which the time interval between neighbouring points is TR*Nfast, where Nfast is the number of k-space indexes along the fast encoding direction. The pulse diagram of the ISME GRE sequence is illustrated in figure 1E. With ISME, the shift of the k-space index is implemented by means of short gradient pulses (‘*blips*’) along the slow phase-encode direction, as in Echo Planar Imaging (EPI).^50^ However, unlike EPI, the data from the consecutive readouts of the multi-echo train are stored separately to obtain an individual image at each echo time. To reduce acquisition time, appropriate modulation of the amplitude of the blip gradients allows undersampling of k-space according to e.g. the GRAPPA^51^ or CAIPIRINHA^52^ filling patterns used for Echo Planar Time-resolved Imaging (EPTI)^53,54^ or segmented EPI.^55,56^

With ISME, the multi-echo data at a single k-space index is acquired over a duration TR*Nfast*Necho, where Necho is the number of echoes. To fully populate the k-TE space of the data with the diagonal sampling pattern of ISME (top-left and bottom right corners of Figure 1C), the sampling trajectory of ISME is initiated beyond the k-space boundary determined from the image encoding parameters, at k=-(Necho-1) and k=kend-(Necho-1). The RF receivers are blanked in this region of k-space (shaded areas of Figure 1C), resulting in an increase of scan time by TR*Nfast*(Necho-1).

### 2.2. MRI data acquisition

Eleven healthy volunteers (4 females, mean age=33±9 years old) were scanned on a 3T MRI system (Magnetom Prisma; Siemens Medical Systems, Erlangen, Germany) using a 64-channel head coil. Written informed consent was obtained from each participant prior to participation and the study was approved by the local ethics committee. The data from one participant was excluded from analysis due to strong motion artifacts (Supplementary material 1). Acquisition of the GRE data was conducted using a custom-made multi-echo 3D FLASH pulse sequence. The k-space trajectory was linear Cartesian. The main acquisition parameters are highlighted in Table 1. 14 echo images were acquired with a bipolar readout, with echo times TE=1.25ms to 16.85ms with 1.2ms echo spacing (protocol 1). The repetition time was TR=21ms, the RF excitation angle 12°, the voxel size was 1.2x1.2x1.2mm^3^ and the field of view 208×192×144mm^3^. The images were reconstructed on the MRI scanner with adaptive coil combination^57,58^ and GRAPPA^51^ with an acceleration factor of 2. Phase navigators were used to correct for systematic phase differences between positive and negative readouts.^59^ In pilot experiments, ISME was observed to mitigate aliasing artefacts of non-cardiac origin such as those due to eye movement (Supplementary material 2). To keep the focus of this study specifically on cardiac-induced effects, the imaging volume was tilted by 30 degrees in the sagittal plane to displace this artefact below the brain. The total scan time was 5:29 minutes for the standard multi-echo trajectory and 6:08 minutes (+12%) for ISME due to the k-space boundary excursions. In experiment 1, each sampling strategy was acquired 3 times, in a randomized order, to quantify the reproducibility of the data across repetitions. In experiment 2, one of the participants (male, 31 years old) underwent an additional MRI exam with protocol 1. To assess the incoherent averaging of cardiac-induced signal instabilities across multiple repetitions of the standard multi-echo trajectory, experiment 2 involved 9 repetitions of data acquisition with the standard multi-echo trajectory and 3 repetitions of data acquisition with ISME.

**Table 1:** Scanning protocols.

|  | TR | TE | Resolution | Tilt | Duration | # repetitions |
| --- | --- | --- | --- | --- | --- | --- |
| Protocol 1 | 21ms | $TE_{\min}=1.25\text{ms}$;<br>echo spacing = 1.2ms | $(1.2 \text{ mm})^3$ | $30^\circ$ | 5:29<br>(Standard)<br>6:08 (ISME) | Experiment 1: 3<br>(Standard); 3 (ISME)<br>Experiment 2: 9<br>(Standard); 3 (ISME) |
| Protocol 2 | 35.55ms | $TE_{\min}=2.3\text{ms}$;<br>echo spacing = 2.14ms<br>spacing | $(0.8 \text{ mm})^3$ | $0^\circ$ | 18:52<br>(Standard)<br>20:28 (ISME) | 1 |

One of the participants (male, 44 years old) underwent an additional MRI exam at a higher resolution of 0.8x0.8x0.8mm^3^ (protocol 2, see Table 1; other parameters the same as above). No tilt of the imaging volume was used). During this exam, one multi-echo dataset with the standard multi-echo trajectory (18:52 minutes) and one with the proposed ISME (20:28 minutes, +8%) were acquired. This data is available online (10.5281/zenodo.13364051).

To allow accurate tissue segmentation and delineation of grey matter regions, a multi-parameter relaxometry protocol was also used for the acquisition of multi-echo 3D FLASH data with magnetization transfer (MT), proton density (PD) and T1-weighted contrast (RF excitation angle = 6°, 6° and 21° respectively; TR/TE=24.5/2.34ms; voxel size: 1.5x1.5x1.5mm^3^; field of view: 176×240×256mm^3^). 8 echo images were acquired for the T1- and PD-weighted contrasts and 6 for the MT-weighted contrast. B1- field mapping data (voxel size: 4x4x4mm^3^; TR/TE=200/39.1ms) and B0-field mapping data (voxel size: 3x3x2.5mm^3^; TR=700ms; TE1/TE2=4.92/7.38ms) were also acquired.^60,61^

### 2.3. Image segmentation

Maps of MTsat^62^ were computed from the MT-, PD- and T1-weighted images using the hMRI toolbox (https://hMRI.info).^63^ The MTsat maps and GRE data acquired with the standard multi-echo trajectory and ISME were co-registered using Statistical Parametric Mapping (SPM12, Wellcome Centre for Human Neuroimaging, London, UK).

Grey and white matter probabilities maps were obtained from the segmentation of the MTsat maps with SPM Unified Segmentation.^64–66^ Whole-brain masks were defined from the voxels with a grey or white matter probability above 0.95. Because B0-field inhomogeneities affect the transverse decay of the MR signal,^67^ voxels in the proximity of the air/tissue interface (e.g. orbitofrontal cortex, amygdala, temporal lobe, hippocampus) were removed from the masks. Brainstem and cerebellum regions of interest (ROIs), in close proximity to the ventricles, carotid arteries and large blood vessels were defined from the labelled data provided by Neuromorphometrics, Inc. (http://neuromorphometrics.com/) under academic subscription, and were set to contain voxels with a grey or white matter probability above 0.95. Additional occipital, frontal, parietal and temporal grey matter ROIs were also computed from the Neuromorphometrics atlas. White matter ROIs of the corticospinal tract, inferior longitudinal fasciculus and optic radiation were computed from the JHU DTI-based atlases (https://identifiers.org/neurovault.collection:264).^68–70^

### 2.4. Relaxometry

Data were analysed using bespoke analysis scripts written in MATLAB R2021a (The Mathworks, Natick, MA). Before the estimation of R2*, the distribution of signal intensities in the background voxels of the magnitude images was fitted with a Rician distribution. The value of the noncentrality parameter was deducted from the signal intensities to suppress the noise floor in these images.^71–73^ The signal intensities in each voxel of the magnitude images were then fitted with the following model:

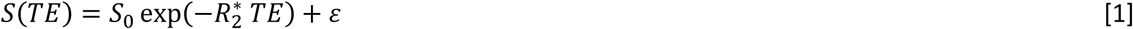

where 𝑆_0_ is the signal amplitude at 𝑇𝐸 = 0 and 𝜀 is the residual error. Fitting was conducted using the non-linear least square (NLS) *lsqnonlin* MATLAB function with a trust-region-reflective algorithm. R2* was bounded between 1 to 80s^-1^ and its initial value was 20s^-1^. The root-mean-squared error (RMSE) of the fit was computed as a measure of the lack of fit. To investigate the effect of the number of echoes, R2* estimation was conducted using all 14 acquired echo data, 7 echoes (echo numbers 1, 3, 5, 7, 9, 11 and 13), 5 echoes (echo numbers 1, 4, 7, 10 and 13), 4 echoes (echo numbers 1, 5, 9, 13) and 3 echoes (echo numbers 1, 6 and 11).

The NLS approach for the estimation of R2* assigns an equal contribution of all echo images to the error estimates, regardless of their respective echo times. Alternatively, we also computed maps of R2* using a nonlinear weighted least squares (NWLS) approach, with weights that reflect the noise level in each individual echo image. Estimation of the image-specific weights was conducted on the model of the QUIQI technique, which addresses the effect of motion degradation on the noise level in image analyses.^74–76^ The noise level in the multi-echo data was modelled as a polynomial function of the echo time. The polynomial coefficients were estimated using the implementation of Restricted Maximum Likelihood (ReML) in SPM^77^ (Supplementary material 3, Matlab code available online (10.5281/zenodo.14808609)). The optimal order of this polynomial, which maximized the estimates of the evidence lower bound (ELBO) provided by ReML, was 4. The image-specific weights were computed as the inverse of the noise covariance matrix estimated by ReML with the optimal noise model. These weights were then used to estimate R2* according to eq. [1], using the non-linear least square *lsqnonlin* MATLAB function.

### 2.5. Quantitative susceptibility mapping

Quantitative magnetic susceptibility maps were generated from the phase of the MR data using bespoke scripts adapted from https://github.com/fil-physics/MPM_QSM. We used ROMEO^78^ for the unwrapping of the phase images and ASPIRE^59^ to remove the systematic phase differences between odd and even echo images. After segmentation of the first echo magnitude image with SPM’s Unified Segmentation,^66^ brain masks were defined to include voxels with a combined grey matter, white matter and CSF probability above 0 and with values of the ROMEO^78^ phase quality metric above 0.3. Any holes present within the resulting mask were subsequently filled using the MATLAB function *imfill*. The spurious contribution of the background field to the phase data was removed using the Projection onto Dipole Fields algorithm^79^ available in the SEPIA toolbox.^80^ Finally, dipole inversion was conducted using the STAR-QSM algorithm^81^ available in the SEPIA toolbox, using the entire brain as a reference.

To investigate the effect of the number of echoes, QSM estimation was conducted using all 14 acquired echo data, 7 echoes (echo numbers 1, 3, 5, 7, 9, 11 and 13), 5 echoes (echo numbers 1, 4, 7, 10 and 13), 4 echoes (echo numbers 1, 5, 9, 13) and 3 echoes (echo numbers 1, 6 and 11).

### 2.6. Statistical analyses

The effects of cardiac pulsation on the MR signal lead to changes of the apparent R2* across the cardiac cycle.^18^ With standard multi-echo trajectories, the multi-echo data at a given k-space frequency is acquired for a given phase of the cardiac cycle and cardiac-induced signal instabilities increases the variability of the data across repetitions.^18^ With ISME, multi-echo data representative of the whole cardiac cycle are acquired for each repetition (Figure 1). We conducted statistical analyses to test if this leads to reduced data variability compared to standard multi-echo trajectories. Regional estimates of the standard deviation (SD) of R2* and 𝜒 across repetitions were computed as the average in each ROI of the voxel-wise SD values. We conducted paired Student’s t-tests of the regional SD estimates, for data acquired using the standard multi-echo trajectory and ISME.

With ISME, the effect of cardiac pulsation on signal decay does not follow the exponential dependence on echo time of data acquired with standard multi-echo trajectories. We conducted statistical analyses to verify that this leads to higher levels of RMSE. Regional estimates of the RMSE were computed as the average in each ROI of the voxel-wise RMSE values obtained from the estimation of R2*. We conducted paired Student’s t-test analyses of the regional RMSE levels, for data acquired using the standard multi-echo trajectory and ISME.

## 3. Results

In individual echo images acquired using both the standard multi-echo trajectory and ISME, the k-space data along the slow phase encoding direction are acquired at different phases of the cardiac cycle, leading to spatial aliasing of pulsating veins (figure 2A and 2C, data acquired with protocol 2). With the standard multi-echo trajectory, the distribution of the cardiac phase along the slow phase encoding direction is unchanged across echo images (Figure 1A) and the resulting R2* maps also display spatial aliasing (Figure 2B). With ISME, the distribution of the phase of the cardiac phase along the slow phase encoding direction differs between echo images (Figure 1C) and the R2* maps computed from the signal decay across echoes show a reduced level of aliasing (Figure 2D).

**Figure 2:**
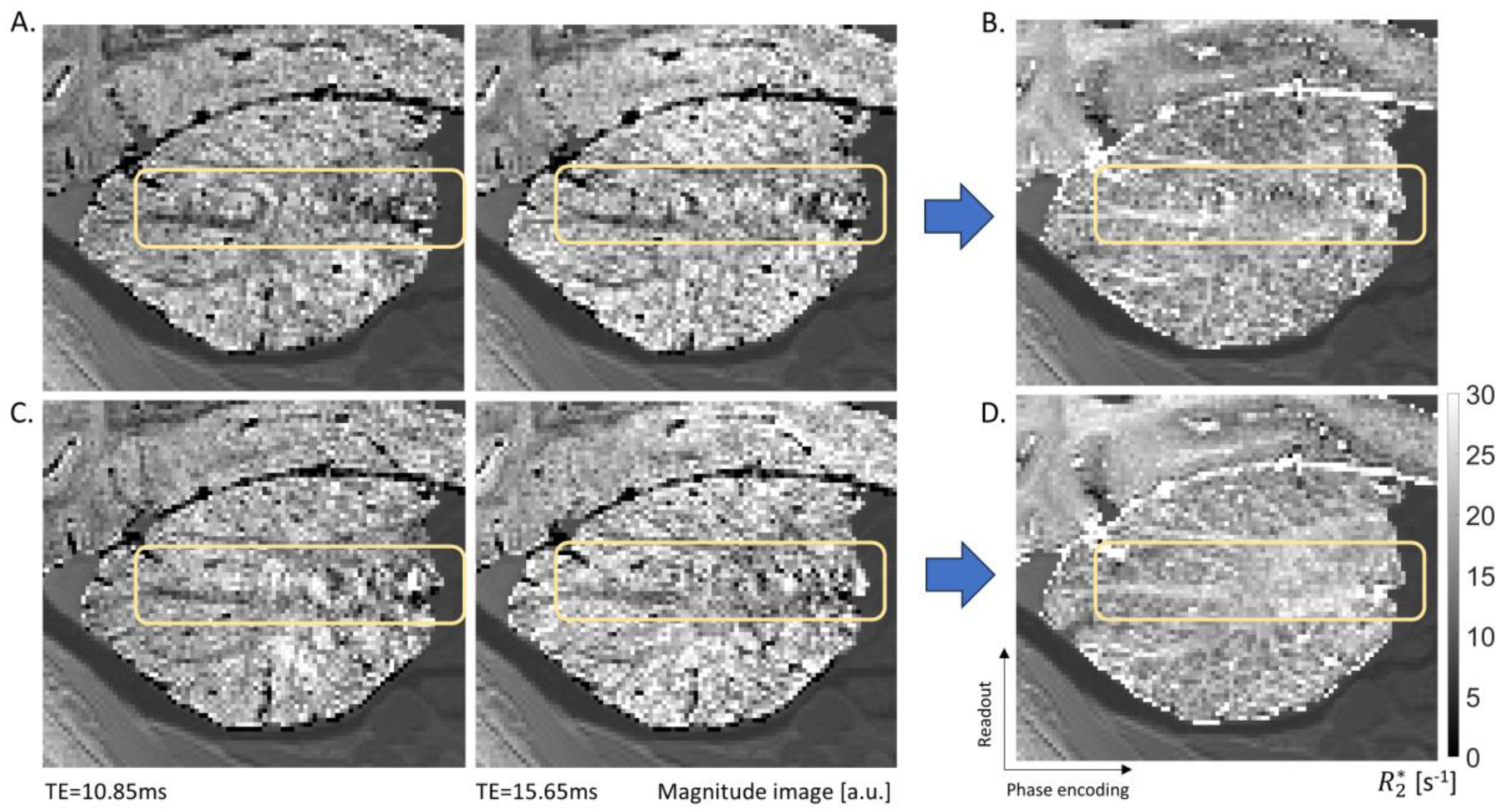
Example GRE images and corresponding R2* maps (scanning protocol 2). (A) GRE images acquired with the standard multi-echo trajectory (TE = 10.85 ms and 15.65ms). (B) Corresponding R2* map computed from the signal decay across echoes. (C) GRE images acquired with ISME (TE = 10.85 ms and 15.65ms). (D) Corresponding R2* map computed from the signal decay across echoes. The light yellow rectangles highlight aliasing of pulsating signal originating from the nearby carotid artery, visible on the individual GRE images and the R2* map computed from the data acquired with the standard multi-echo trajectory. This aliasing is not apparent in the R2* map computed from the data acquired with ISME.

The SD of R2* across repetitions can reach up to 5s^-1^ in inferior brain regions in data acquired with the standard multi-echo trajectory (Figure 3A). The most affected areas include the cerebellum and brainstems – two of the regions most affected by cardiac-induced signal instabilities.^18^ The SD of R2* is much reduced in inferior brain regions with the ISME approach, leading to variability maps that are more spatially uniform. With ISME, the SD of R2* across repetitions is reduced by 25, 26, and 21% in the cerebellum, brainstem and whole brain compared to the standard multi-echo trajectory (Figure 3B, p≤0.001). Regional estimates of the standard deviation of R2* across repetitions are provided in Table 2. The SD of R2* across repetitions decreases with an increasing number of echo images used in the estimation of R2* (𝑁_𝑒𝑐ℎ𝑜𝑒𝑠_) (Figure 3C). The sharpest decrease takes place for 𝑁_𝑒𝑐ℎ𝑜𝑒𝑠_ ≲ 7 and is about twice as large with ISME than with the standard multi-echo trajectory. With the standard multi-echo trajectory, the SD of R2* maps computed from *Nav*=2 repetitions is comparable to that of ISME with *Nav*=1 repetition, i.e. a 100% scan time increase (Figure 3D).

**Figure 3:**
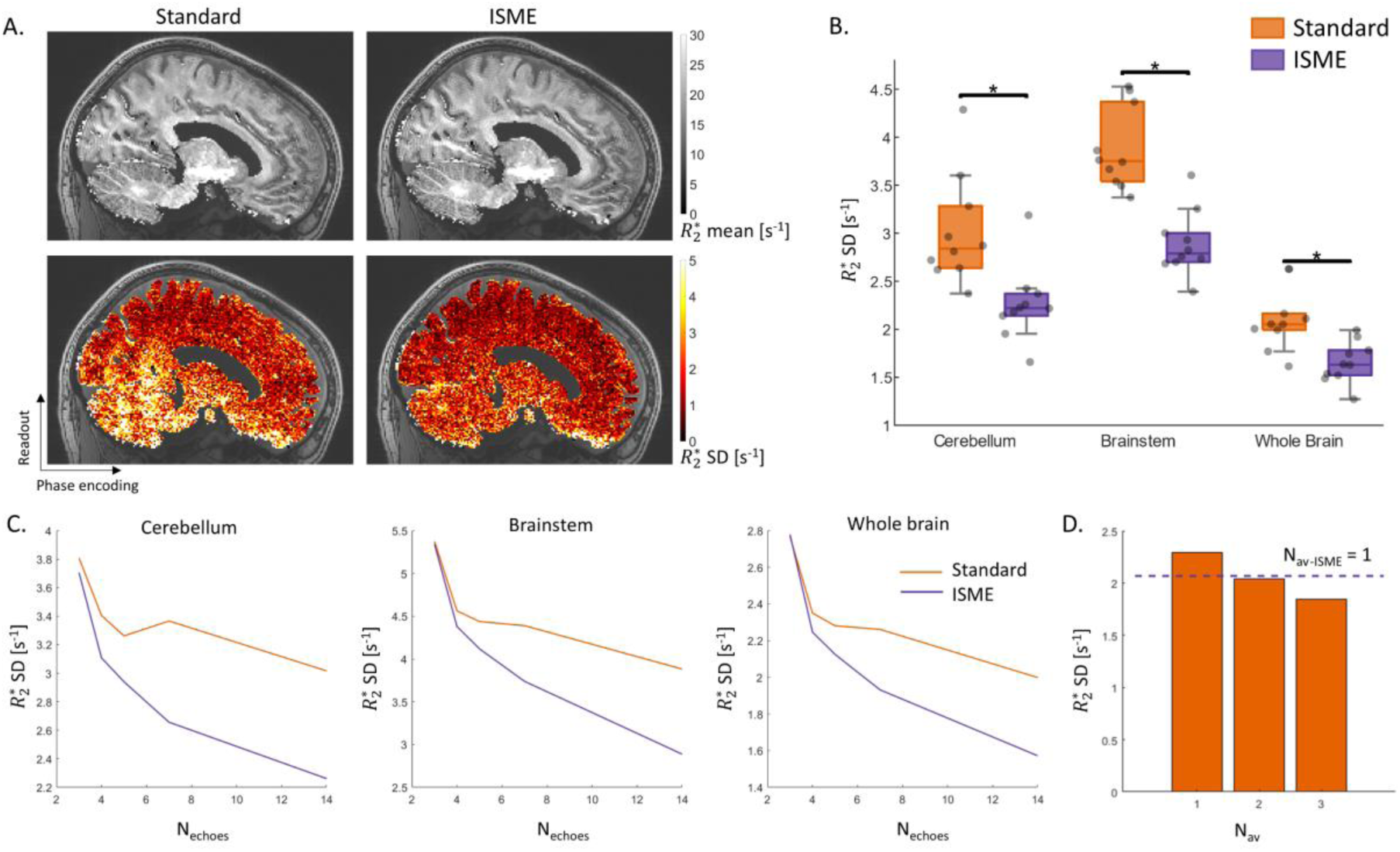
Mean and SD of R2* across repetitions (scanning protocol 1). (A) Example mean and SD across repetitions of R2* maps computed from data acquired with the standard multi-echo trajectory and ISME. (B) ROI-averaged SD of R2* in the cerebellum, brainstem and whole brain for each subject, computed from data acquired with the standard multi-echo trajectory and ISME. (C) Whole-brain average of the SD across repetitions of R2* maps computed from an increasing number of echo images (Nechoes). (D) Average SD across repetitions of R2* maps computed from Nav=1, 2, and 3 datasets acquired with the standard multi-echo trajectory and from Nav-ISME=1 dataset acquired with ISME (Nechoes=14). (A-C) were computed from the data acquired in experiment 1. (D) was computed from the data acquired in experiment 2.

**Table 2:** Variability of R2* estimates across repetitions, lack of fit of the R2* estimates (RMSE) and variability of magnetic susceptibility (𝜒) estimates across repetitions, from data acquired with a standard multi-echo trajectories and with ISME. For the data acquired with ISME, the R2* variability and RMSE estimates were computed using nonlinear least squares (NLS) and nonlinear weighted least squares (NWLS).

| | $R_2^*$ SD [$s^{-1}$] | | | RMSE [a.u.] | | | $\chi$ SD [ppb] | |
| --- | --- | --- | --- | --- | --- | --- | --- | --- |
|  | Standard | ISME NLS | ISME NWLS | Standard | ISME NLS | ISME NWLS | Standard | ISME |
| Brainstem | $3.02 \pm 0.57$ | $2.26 \pm 0.39$ | $2.20 \pm 0.35$ | $2.85 \pm 0.30$ | $3.34 \pm 0.48$ | $3.27 \pm 0.43$ | $12.63 \pm 2.32$ | $9.66 \pm 1.52$ |
| Cerebellum | $3.88 \pm 0.43$ | $2.89 \pm 0.34$ | $2.83 \pm 0.29$ | $2.54 \pm 0.17$ | $2.86 \pm 0.20$ | $2.77 \pm 0.18$ | $10.41 \pm 1.54$ | $7.09 \pm 1.12$ |
| Whole brain | $2.00 \pm 0.31$ | $1.57 \pm 0.20$ | $1.56 \pm 0.20$ | $2.58 \pm 0.19$ | $2.97 \pm 0.36$ | $2.95 \pm 0.34$ | $8.72 \pm 1.67$ | $6.57 \pm 1.31$ |
| Occipital lobe | $1.99 \pm 0.40$ | $1.64 \pm 0.30$ | $1.60 \pm 0.29$ | $3.03 \pm 0.26$ | $3.42 \pm 0.43$ | $3.41 \pm 0.37$ | $12.52 \pm 3.53$ | $10.23 \pm 2.67$ |
| Frontal lobe | $1.77 \pm 0.38$ | $1.46 \pm 0.28$ | $1.47 \pm 0.28$ | $2.70 \pm 0.24$ | $3.17 \pm 0.50$ | $3.19 \pm 0.50$ | $9.99 \pm 1.95$ | $7.87 \pm 2.11$ |
| Parietal lobe | $1.65 \pm 0.27$ | $1.39 \pm 0.21$ | $1.39 \pm 0.21$ | $2.70 \pm 0.17$ | $3.08 \pm 0.34$ | $3.12 \pm 0.32$ | $10.07 \pm 2.25$ | $8.05 \pm 1.51$ |
| Temporal lobe | $2.24 \pm 0.35$ | $1.78 \pm 0.27$ | $1.73 \pm 0.26$ | $2.63 \pm 0.20$ | $3.11 \pm 0.40$ | $3.06 \pm 0.35$ | $9.36 \pm 2.18$ | $6.73 \pm 1.81$ |
| Corticospinal tract | $2.38 \pm 0.28$ | $1.85 \pm 0.22$ | $1.84 \pm 0.20$ | $2.39 \pm 0.12$ | $2.66 \pm 0.24$ | $2.63 \pm 0.22$ | $7.44 \pm 1.29$ | $5.32 \pm 0.97$ |
| Inferior Longitudinal Fasciculus | $2.24 \pm 0.41$ | $1.68 \pm 0.35$ | $1.65 \pm 0.23$ | $2.28 \pm 0.21$ | $2.78 \pm 0.36$ | $2.72 \pm 0.31$ | $7.58 \pm 1.64$ | $5.43 \pm 1.49$ |
| Optic radiation | $1.63 \pm 0.31$ | $1.30 \pm 0.20$ | $1.28 \pm 0.18$ | $2.64 \pm 0.29$ | $2.98 \pm 0.44$ | $2.95 \pm 0.39$ | $6.86 \pm 1.97$ | $5.10 \pm 1.32$ |

Much of these observations are also applicable to maps of 𝜒 (Figure 4A,B). ISME reduces the SD of 𝜒 estimates across repetitions by 24, 32, and 23% in the cerebellum, brainstem, and whole brain, respectively, compared to the standard multi-echo trajectory (Figure 4C, p≤0.001). Regional estimates of the standard deviation of 𝜒 across repetitions are provided in Table 2. The SD of 𝜒 across repetitions decreases with an increasing number of echo images used in the estimation of 𝜒. This decrease is comparable with ISME and with the standard multi-echo trajectory (Figure 4D). The SD across repetitions of 𝜒 maps computed from *Nav*=2 and *Nav*=3 datasets acquired with the standard multi-echo trajectory decreases by ∼30% and ∼36% respectively (Figure 4E).

**Figure 4:**
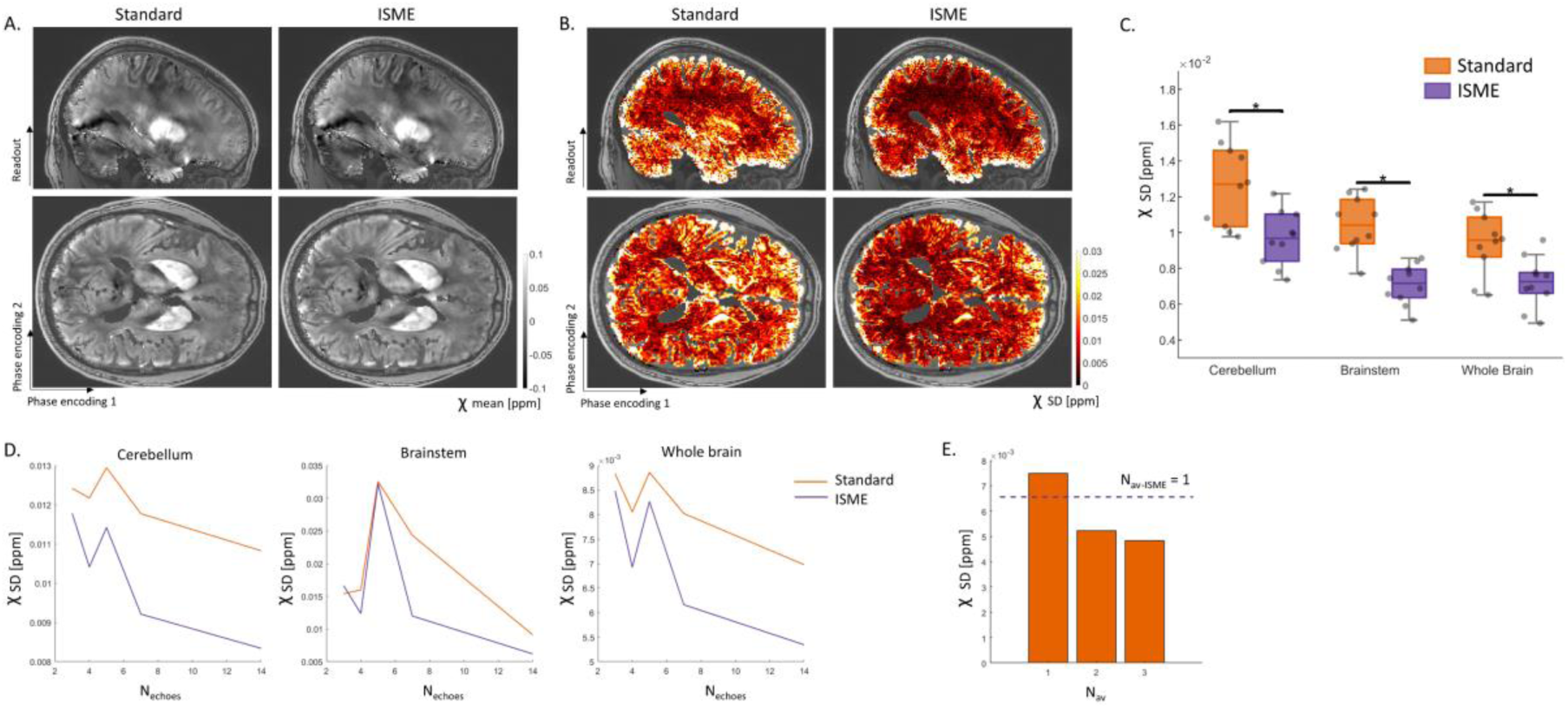
Mean and SD of magnetic susceptibility (𝜒) across repetitions (scanning protocol 1). Example mean (A) and SD (B) across repetitions of 𝜒 maps computed from data acquired with the standard multi-echo trajectory and ISME. (C) ROI-averaged SD of 𝜒 estimates in the cerebellum, brainstem and whole brain for each subject, computed from data acquired with the standard multi- echo trajectory and ISME. (D) Whole-brain average of the SD across repetitions of 𝜒 maps computed from an increasing number of echo images (Nechoes). (E) Average SD across repetitions of 𝜒 maps computed from Nav=1, 2, and 3 datasets acquired with the standard multi-echo trajectory and from Nav-ISME=1 dataset acquired with ISME (Nechoes=14). (A-D) were computed from the data acquired in experiment 1. (E) was computed from the data acquired in experiment 2.

With the standard multi-echo trajectory, the maps of the RMSE between the multi-echo data and the exponential fits show a sharp decrease from the brain periphery to central brain regions that mirrors the sensitivity profile of the receive coil (Figure 5A). Data acquired with the proposed ISME approach exhibit higher RMSE levels and a high level of spatial aliasing along the slow phase encoding direction. This increase is stronger in inferior brain areas and around the circle of Willis: the RMSE increases by 18, 13 and 15% in the cerebellum, brainstem and whole brain with the ISME approach (Table 2 and Figure 5B, p≤0.001).

**Figure 5:**
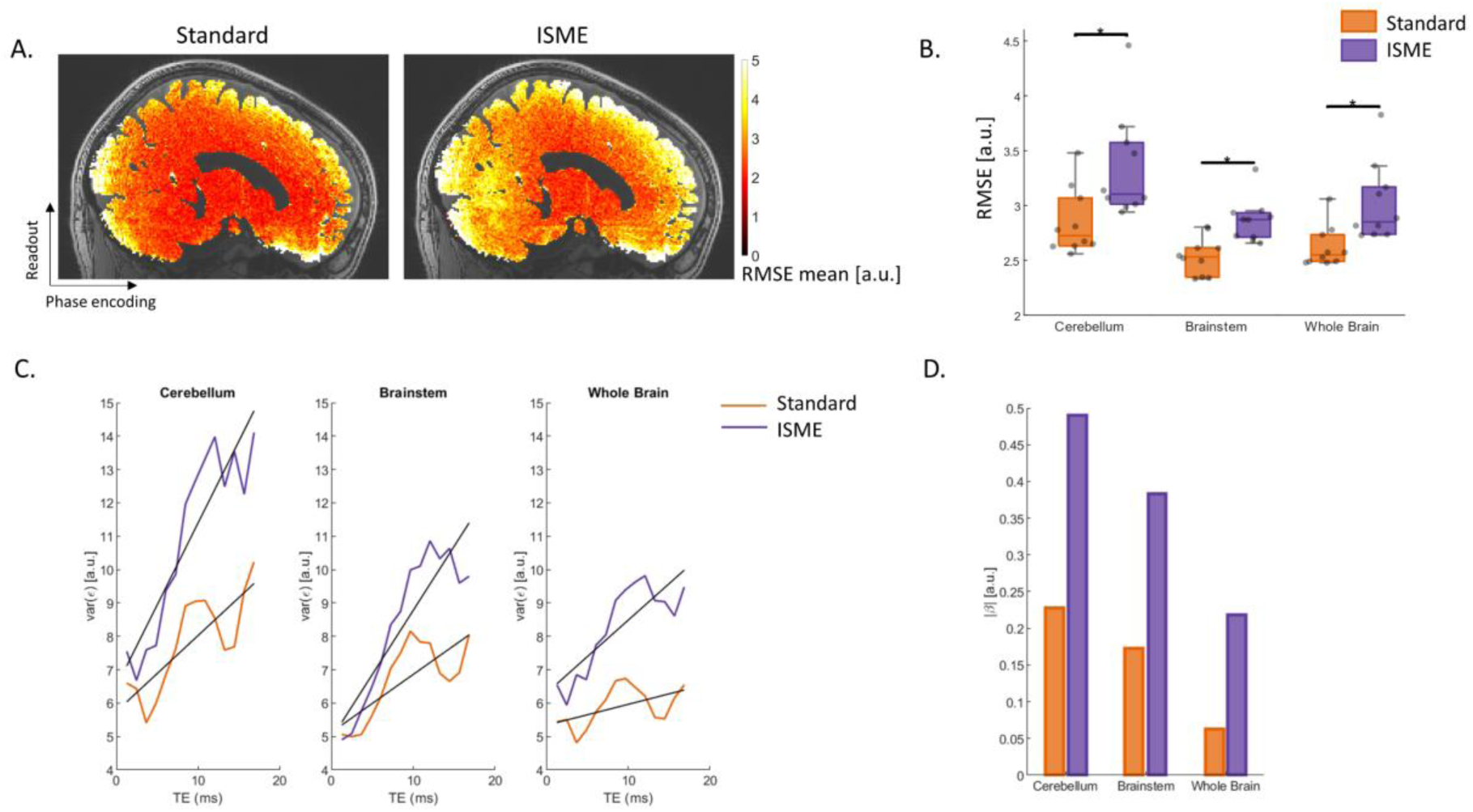
Lack of fit of the R2* estimates (scanning protocol 1, experiment 1). (A) Example map of the mean root mean square error (RMSE) of R2* fits across repetitions, computed from data acquired with a standard multi-echo trajectory and with ISME. (B) Regional RMSE levels in the cerebellum, brainstem and whole brain. (C) Variance of the residuals (ε) across repetitions as a function of echo time for both the standard multi-echo trajectory (orange) and ISME (purple). (D) Rate of increase of the variance of the residuals with the echo time.

The variance across repetitions of the residual error increases with echo time for both acquisition strategies (Figure 5C). However, this increase is more pronounced with ISME: assuming a linear increase, the slope reaches 0.49, 0.38 and 0.22 [a.u./ms] in the cerebellum, brainstem and whole brain with ISME and 0.23, 0.17 and 0.06 [a.u./ms] with the standard multi-echo trajectory in these regions (Figure 5D).

The proposed NWLS approach computes weights that are lower at long echo times due to the higher noise level in the corresponding echo images (Figure 6A). Using the NWLS approach to estimate R2* improves the distribution of the residuals across echo times in data acquired with ISME. However a coherent pattern, with higher residuals at intermediate echo times, remain (Figure 6B). Assuming a linear increase of the residual level with echo time, the slope reduces to 0.05, 0.03 and 0.19 [a.u./ms] in the cerebellum, brainstem and whole brain (0.49, 0.38 and 0.22 [a.u./ms] with NLS, see above) (Figure 6C). The proposed NWLS approach further reduces the SD of R2* by 3% in the cerebellum, 2% in the brainstem and 1% in the whole-brain compared to the standard NLS (Table 2, p≤0.001).

**Figure 6:**
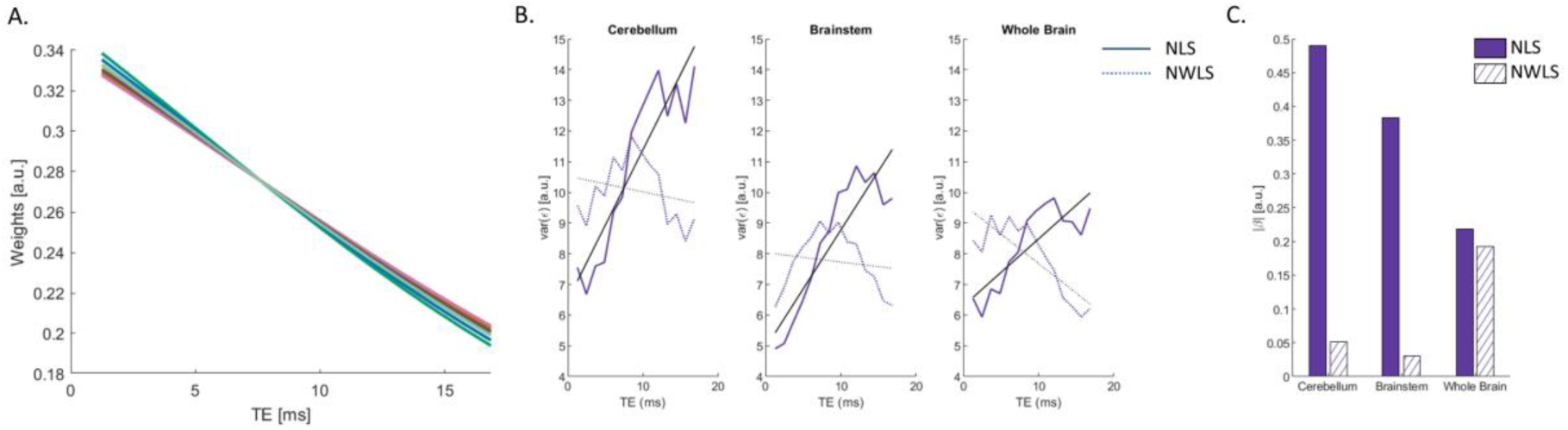
Lack of fit of the R2* estimates with the proposed NWLS approach (scanning protocol 1, experiment 1). (A) Weights computed with the proposed NWLS approach for each study participant. (B) Variance of the residuals (ε) across repetitions as a function of echo time, following estimation of R2* from the ISME data with the NLS and the NWLS approaches (B). (C) Rate of increase of the variance of the residuals with the echo time with the NLS and the NWLS approaches.

## 4. Discussion

Cardiac pulsation generates exponential-like effects on the transverse decay of multi-echo GRE data, leading to systematic changes of R2* across the cardiac cycle^18^. With standard trajectories, the multi-echo data is acquired at the same phase of the cardiac cycle, for each k-space frequency, leading to a bias on the R2* estimates. Because the effect of cardiac pulsation depends on the timing of data acquisition relative to the cardiac cycle, cardiac-induced signal instabilities increase the variability of brain maps of the transverse relaxation rate (R2*) and quantitative susceptibility (QSM) across repetitions. In this work, we proposed a new sampling strategy termed Incoherent Sampling of Multi-Echo data (ISME), based on shifting k-space location between each readout of the multi-echo train. With ISME, the multi-echo data of each k-space frequency is made up of points acquired at different phases of the cardiac cycle, mitigating the effects of cardiac pulsation across echoes.

With ISME, estimation of R2* and 𝜒 effectively averages out the effect of cardiac pulsation and leads to improved repeatability of the data across repetitions: over the whole brain, ISME reduces the variability of R2* across repetitions by 21% compared to standard multi-echo trajectories. Similarly, the variability of 𝜒 estimates is reduced by 23% with ISME. The reduction in variability were most pronounced in inferior brain regions such as the brainstem and cerebellum (25-26% for R2* maps and 24-32% for 𝜒 maps). With standard multi-echo trajectories, a comparable reduction in variability requires the computation of R2* maps across 2 averages and a 100% increase in scan time. The SD of R2* and 𝜒 across repetitions decreases when the number of echo images used in their estimation increases. With ISME, the sharper decrease of the variability of R2* is consistent with the improved sampling of cardiac-induced signal instabilities across echo images. Around 7 echo images are required in the multi-echo data to achieve near-optimal reduction of the variability of R2* across repetitions. Maps of the SD of R2* and 𝜒 estimates across repetitions are spatially more uniform with ISME. This reduction of the variability of R2* and 𝜒 maps is similar to that from an alternative sampling strategy based on optimizing the number of samples at each k-space location based on the local amplitude of cardiac-induced signal instabilities.^49^ The scan time increase is also comparable between both methods (12% vs 14%). However for ISME, this scan time increase is due to the diagonal pattern used to traverse the k-TE space of the data, which imposes a k-space boundary beyond that determined from the image encoding parameters. The use of advanced trajectories such as Gilbert curves might alleviate any scan time increase.^47,48^ Also, ISME acts on all k-space frequencies of the data and might better mitigate spatially localized effects of cardiac pulsation.

With both standard multi-echo trajectories and ISME, neighbouring k-space data are acquired at different phases of the cardiac cycle. Therefore, individual echo images display spatial aliasing of pulsating MR signal across the field of view of the image. However, for the standard multi-echo trajectory, this aliasing is unchanged between echo images and the R2* maps computed from the multi-echo data also exhibit aliasing. With the proposed ISME approach, aliasing artifacts vary across individual echo images because the phase of the cardiac cycle between neighbouring k-space indexes varies between images. As a result, spatial aliasing of pulsating MRI signal arising from e.g. arterial vessels is strongly reduced in R2* maps. In the venous system, blood flow is largely constant and a dedicated flow-compensation technique should be preferred.^15^ We highlight that ISME can be combined with flow-compensation techniques^15,82^ or navigator-based corrections of breathing effects,^83,84^ which were not used in this study.

With standard multi-echo trajectories, cardiac pulsation induces exponential-like effects on the decay of the multi-echo data that depends on the phase of cardiac cycle at the time of acquisition of the data. ISME allows the incoherent sampling of cardiac-induced signal instabilities in the multi-echo data, leading to a 13-18% increase in RMSE compared to standard multi-echo trajectories. Note that the higher RMSE with ISME does not imply that the effective noise level is higher: with ISME, the RMSE measures also include cardiac-induced effects that only manifest in the variability of R2* and 𝜒 maps with standard multi-echo trajectories. ISME also leads to a more pronounced increase of the residuals with increasing echo time. To address this, we introduced a novel nonlinear weighted least squares (NWLS) approach for the estimation of R2*, in which the weights are computed from the noise level of individual echo images using restricted maximum-likelihood (ReML), a statistical method that allows the modelling of noise heteroscedasticity within a dataset. ReML has found multiple applications in neuroimaging, in which noise has different physiological origins, such as the modelling of time correlations in functional MRI time series,^77^ the modelling of motion-induced effects^74,85^ or of non-uniform error variances in group-level analyses.^86^ The proposed NWLS approach improves the distribution of the residuals across echoes, although a coherent pattern remains, and brings in an additional small reduction in variability of R2* across repetitions (1-3%). This approach may bring more pronounced benefits with more complex models of transverse relaxation,^73,87^ longer echo times or higher field strengths (e.g. 7T). Combined, the proposed ISME approach and NWLS estimation might allow the data to better comply with the requirements of independent and identically distributed noise for the denoising of multi-echo data.^88,89^ Here, analysis of the RMSE and estimation using the proposed weighted least square approach was not conducted for 𝜒 as we only had access to compiled code for the estimation for this MRI parameter.

The proposed ISME method is a passive mitigation strategy which in effect averages out physiology- induced signal instabilities across the multiple echoes of the GRE data. Breathing induces periodic fluctuations of the main magnetic field B0 in the head due to changes of the air volume in the lungs.^16,35,90,91^ The multi-echo data is acquired for different phases of the breathing cycle as well as cardiac cycle. Therefore, ISME acts on instabilities that arise from both cardiac pulsation^18^ and breathing,^16,83,84^ as long as these instabilities exhibit a semi-periodic behaviour.^47^ The 20-25% reduction of R2* variability with ISME is lower than recent estimates of cardiac-induced R2* change (∼35%),^18^ presumably due to the higher image resolution. While this study drew its motivation from the effect of cardiac pulsation alone, it is likely that the reduction of R2* and 𝜒 variability with ISME also arises from the mitigation of breathing-induced effects. Breathing has been identified as a dominant source of signal instabilities, albeit at longer echo times than the present study.^16,83,84^

The key novel aspect of ISME is the acquisition of multi-echo data at different phases of the cardiac cycle. The benefits of ISME were demonstrated using a simple GRE sequence as a starting point, for which the multi-echo data are acquired at a single k-space frequency after each RF excitation. Alternative strategies such as EPTI^53,54^ or segmented EPI^55,56^ have been introduced that allow dramatic scan time reductions and maintain a dense sampling of the MRI signal across echo times. These strategies divide k-space into segments that are sampled between consecutive shots. With the current implementations of these techniques, the multi-echo data of a given segment is acquired within a single shot. Because the shot-to-shot interval (TR) is typically shorter than the cardiac period, we expect the multi-echo data from one segment to be equally sensitive to cardiac-induced instabilities as the GRE sequence of the current work. Keeping the k-space filling pattern unchanged, implementation of ISME into EPTI or segmented EPI sequences would involve the generalization of spatial scrambling methods^47,48^ to the additional dimension of the echo time, so that the multi-echo data of each segment is acquired across different phases of physiological cycles.

ISME shifts the current value of the k-space index between each readout of the multi-echo train. The k-space index is independent of the physical coordinates of the k-space data, and ISME is readily transferable to most sampling trajectories. Here we implemented ISME in a linear Cartesian trajectory. However, a pseudo-spiral trajectory has been used recently to reduce the level of cardiac-induced signal instabilities by increasing the number of samples near the k-space centre.^49^ The combination of these two strategies could be beneficial e.g. when too few echo images are acquired to effectively sample the cardiac cycle between echoes with ISME alone (𝑁_𝑒𝑐ℎ𝑜𝑒𝑠_ < 7 from the present results). However, hardware limitations (amplitude, slew rate,…) might impose limits on the gradients required to shift the k-space index between the readouts. The impact of the eddy currents induced by these gradients on image quality should be carefully considered and might pose a practical limitation on the implementation of ISME in alternative sampling trajectories.

## 5. Conclusion

In this work, we proposed a new sampling strategy that reduce the effect of cardiac pulsation on estimates of the transverse relaxation rate (R2*) and magnetic susceptibility (𝜒) computed from multi-echo gradient-echo data. The proposed sampling strategy, termed Incoherent Sampling of Multi-Echo data (ISME), is based on shifting the k-space position of the acquired data between each readout of the multi-echo train. With ISME, the multi-echo data is made up of points acquired at different phases of the cardiac cycle. As a result, the variability of R2*/ 𝜒 maps across repetitions is reduced by 20-25% / 23-32% compared to standard multi-echo trajectories. With ISME, the spatial aliasing of pulsating MR signal is incoherent across the set of multi-echo images. This allows effective reduction of this aliasing in the R2* maps computed from the data.

MRI data acquired with ISME show a stronger increase of the residual level with echo time. We introduce a novel nonlinear weighted-least squares (NWLS) approach for the computation of R2* maps, in which the weights are computed from the noise covariance matrix of the data – estimated using Restricted Maximum Likelihood. The proposed NWLS approach improves the distribution of the residuals across echo times.

ISME allows the effective reduction of cardiac-induced signal instabilities in in vivo gradient-echo data and enhances the sensitivity of R2* and 𝜒 estimates to brain change in neuroscience studies.

## Data availability statement

The high-resolution dataset is available online here (10.5281/zenodo.13364051). The Matlab code used for the estimation of the image-specific weights is available online here (10.5281/zenodo.14808609).

## Ethics approval statement

This study received approval from the local Ethics Committee and all participants gave their written informed consent prior to participation.

## 1. Supplementary material

### 1.1. Motion degradation index

The index introduced in Castella et al. ^74–76^ was calculated from each multi-echo dataset to estimate the level of image degradation caused by head motion (‘Motion Degradation Index’, MDI). The distribution of MDI values peaked at MDI ∼3.5s^-1^, reflecting an overall high image quality (Figure S1). However, four datasets from the same subject exhibit MDI values above 4s^-1^, indicative of poor quality due to head motion. The data from this subject was removed from subsequent analyses.

### 1.2. Mitigation of eye movement artefacts

Eye movement during the acquisition of MRI data leads to image aliasing along the slow phase-encoding direction of a 3D-encoded image. When the slow phase-encoding direction is oriented along the anterior-posterior direction of the patient, this aliasing propagates across the brain with a standard multi-echo trajectory (Figure S2A). However, this aliasing artifact is largely mitigated with the proposed ISME approach (Figure S2B). To keep the focus of this study specifically on cardiac-induced effects, the imaging box was tilted by 30 degrees in the sagittal plane to displace this artefact below the brain.^92^

### 1.3. Model optimization for estimation of R2* using NWLS

Fitting a nonlinear model with a weighted LS approach involves two sets of unknowns: the model parameters and the weights, with the latter corresponding to the inverse of the variance of the residuals at each echo time. Estimating both sets of parameters by maximum-likelihood results in biased estimates of (co)variance, which in turn bias the estimated model parameters. In contrast, restricted maximum-likelihood approaches (ReML) yield unbiased estimates of variance. ReML, however, does not lend itself easily to nonlinear models. We therefore devised a two-step procedure, whereby a linear ReML estimate of variance is performed first using a Taylor expansion of the transverse signal decay, and the resulting weights are then used in a nonlinear weighted least square (NWLS) fit. ReML estimation was performed using the function ‘spm_reml’ in SPM (the analysis script is provided with this article). The covariance matrix of the residual noise was assumed diagonal, with weights modelled as a polynomial in TE. As is standard in SPM, we assumed that this covariance matrix is shared across all voxels within a brain volume.

We identified the optimal order of the Taylor expansion (M) from the Akaike information criterion corrected for small sample size (AICc). The AICc rewards the goodness of fit and penalizes the number of estimated parameters.^93^ The minimal values of the AICc were found for Taylor expansion orders of 1 and 2 (Figure S3A). We opted for a Taylor expansion of order M=2 to accommodate local deviations from the linear approximation of the signal decay due to e.g. high decay rate in iron-rich regions (pallidum, red nuclei, substantia nigra,..) or deviations from the exponential behaviour in white matter and sub-cortical areas.^73,87,94^

From the results of the NLS analyses shown in Figure 5C, models of the dependence of the residuals on the echo time of the data were assumed to follow a polynomial form of order N. Model selection was conducted from the estimates of the evidence lower bound (ELBO) provided by the implementation of ReML of the SPM software.^77^ The ELBO favours the reduction of residual errors and penalises model complexity. An optimal noise model maximises the ELBO. Compared to N=0 (uniform residual level across echo times), the optimal noise model was found to be N=3 for data acquired with the standard multi-echo trajectory. The corresponding increase in ELBO was ΔELBO=ELBON=3- ELBON=0=43.3 (Figure S3B). With the proposed ISME method, the optimal polynomial order was N=4. The corresponding increase in ELBO (ΔELBO=ELBON=4-ELBON=0=73.1) is higher than in data acquired with the standard multi-echo trajectory. Image-specific weights were computed as the inverse of the diagonal elements of the noise covariance matrix estimated by ReML. The dependence of the weights on the echo time of the data shows little change for values of N≥4 (Figure S3C).

## Supplementary figures

**Figure S1:**
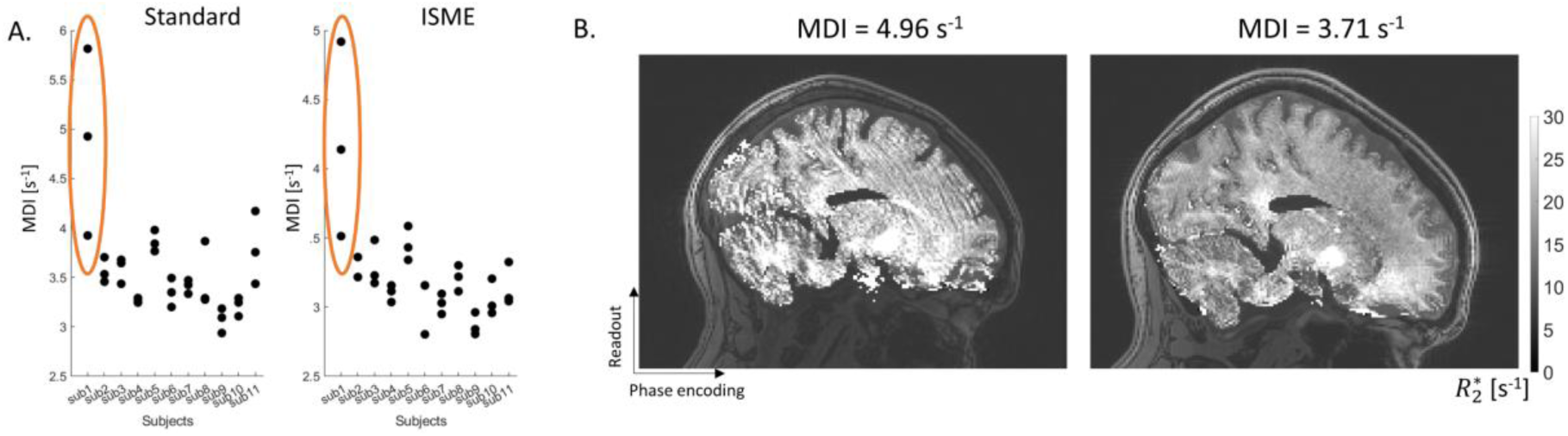
(A) Motion degradation index (MDI) of the datasets acquired with the standard multi-echo trajectory and ISME. The average MDI from subject 1 was 4.5±0.8s^-1^ and the average MDI amongst the other subjects was 3.3±0.3s^-1^. (B) R2* map computed from the first repetition of the standard multi-echo trajectory for subject 1 (MDI=4.96s^-1^) and for one participant with MDI=3.71s^-^

**Figure S2:**
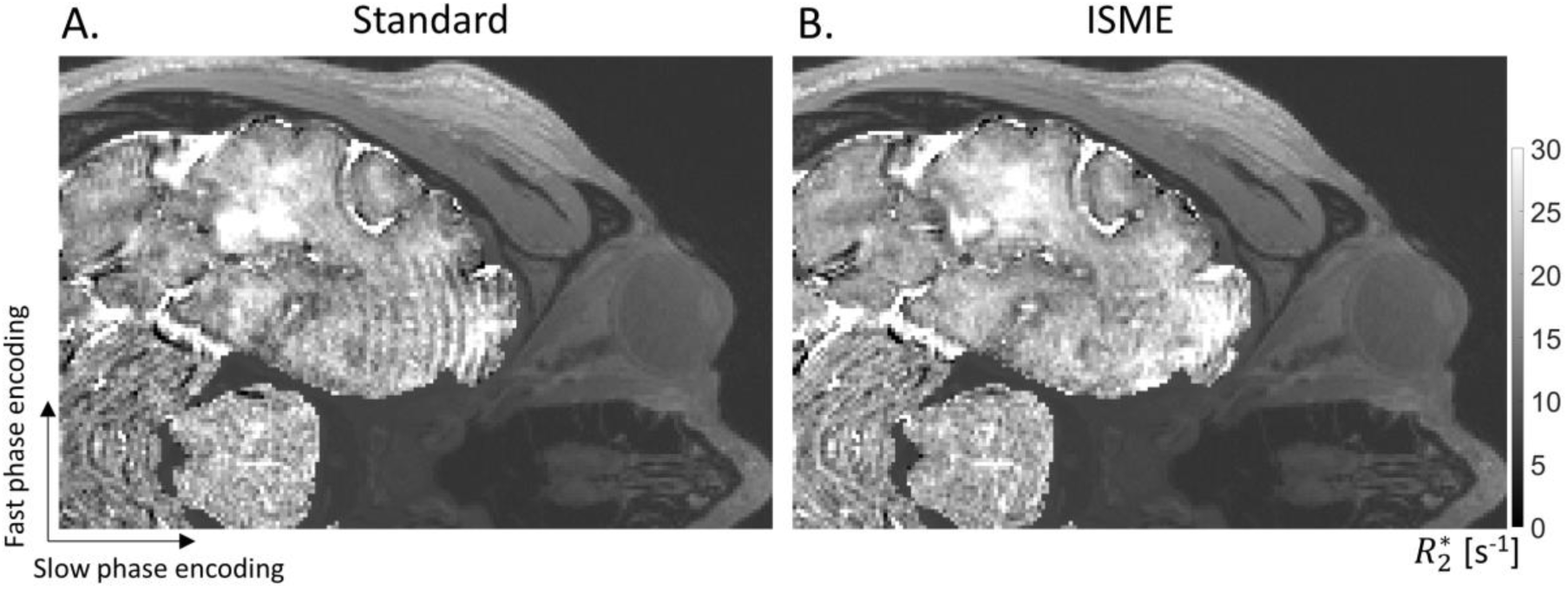
Eye movement during the acquisition of MRI data leads to image aliasing along the slow phase-encoding direction of a 3D-encoded image (blue arrows). This aliasing is present in data acquired with a standard multi-echo trajectory (A) but not with ISME (B).

**Figure S3:**
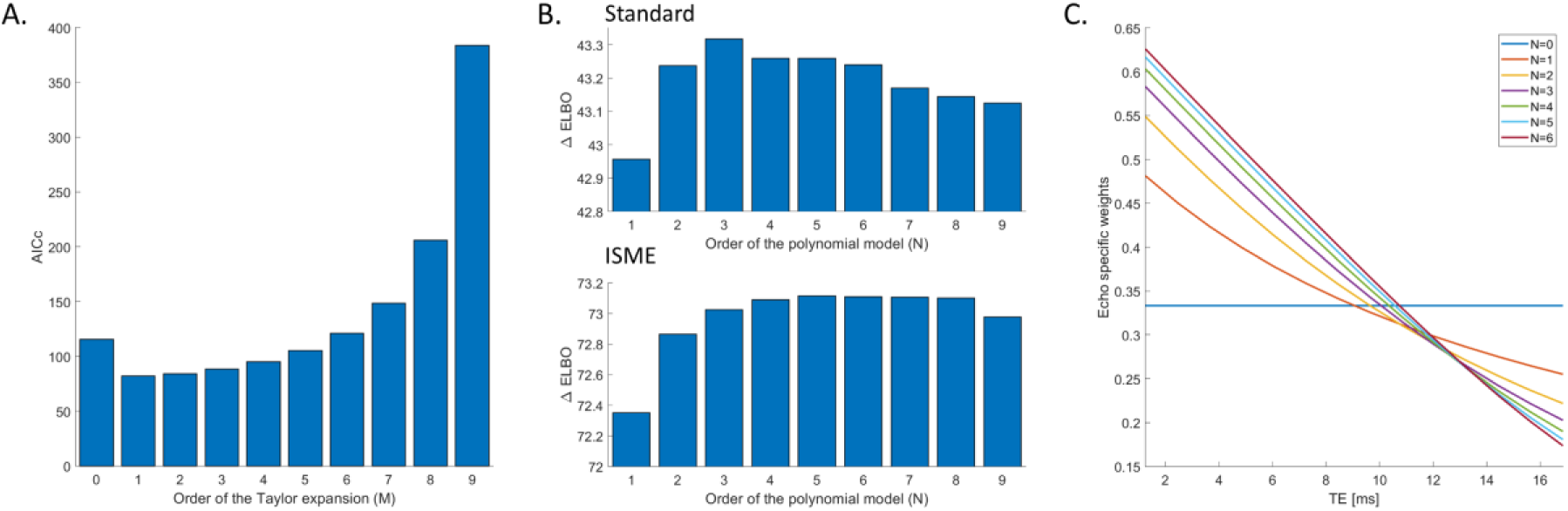
(A) Change in the Akaike information criterion (AICc) with the order of the Taylor expansion of the signal decay (M). (B) Increase of the evidence lower bound (ELBO) with the order of the polynomial model of the dependence of image noise on the echo time of the data (N), compared to N=0 (uniform noise level across echo times), for standard multi-echo trajectory and ISME. (C) Echo specific weights computed for different polynomial orders (M=2).

